# Metagenomic Insights Into Microbial Controls of Carbon Cycling in Alpine Soils

**DOI:** 10.1101/2025.09.22.677713

**Authors:** Kristina Bright, Bence Dienes, Bart van Dongen, Ilya Strashnov, Xingguo Han, Meret Aeppli

## Abstract

Alpine riparian zones span topographic gradients from wet soils on the plain near streams to drier soils on adjacent slopes. These differences in soil moisture are generally associated with shifts in soil redox state from anoxic on the plain to oxic on the slope. In anoxic plain soils, soil organic carbon (SOC) may accumulate due to thermodynamic constraints on microbial activity. Here, we used shotgun metagenomics to examine how microbial diversity and functional potential varies across differing redox conditions on plain and slope soils in two catchments in the Swiss Alps. We complemented these analyses with soil physicochemical characteristics and information on the chemical composition of organic matter. Plain soils had higher SOC stocks and higher relative abundance of phenol compounds relative to slope soils, consistent with SOC preservation and preferential mineralisation of easily degradable organic compounds under anoxic conditions. Microbial communities in plain soils further exhibited greater taxonomic and functional diversity, including an increased potential for anaerobic respiration pathways. Genes for nitrate, iron, and sulfate reduction were linked to *Chloroflexota, Acidobacteria*, and *Desulfobacterota* phyla, respectively. Based on NMDS correlations, electron accepting capacity, calcium content, and pH shaped microbial community composition. Slope soils, by contrast, supported less diverse microbial communities, determined mainly by electron donating capacity and clay content. Our work demonstrates how soil redox conditions and microbial functional potential shape carbon cycling across landscape positions in alpine riparean zones. This mechanistic understanding is critical to anticipate changes in carbon cycling in alpine ecosystems in a changing climate.

## Introduction

In subalpine and alpine ecosystems, more than 90% of ecosystem carbon are stored in soils as a consequence of short plant growing seasons and limitations on the degradation of soil organic matter by microorganisms under harsh climatic conditions (Davidson and Janssens, 2006; Körner, 2021). The fate of organic carbon in the soil is determined by microorganisms that can mineralize organic matter to greenhouse gases or stabilize it within soils (Duan et al., 2023; Gunina and Kuzyakov, 2022). It remains unclear how SOC stocks are linked to microbial community composition and functional potential in (sub)alpine ecosystems.

Alpine riparian zones express strong differences in hydrology and soil biogeochemistry between soils on low-lying plains near streams to those on adjacent slopes (Berhe and Kleber, 2013; Pacific et al., 2011). Plain soils, influenced by shallow groundwater and seasonal water inputs, are periodically saturated, producing oxygen-limited redox conditions where microbial respiration depends on alternative terminal electron acceptors (TEAs). These less energy-efficient pathways slow organic matter decomposition and promote SOC accumulation (Boye et al., 2017; Keiluweit et al., 2016; Schimel and Schaeffer, 2012; Zhang and Furman, 2021). In contrast, slope soils are well drained and maintain more oxidized conditions that support aerobic microbial activity and greater SOC mineralization, resulting in smaller SOC stocks relative to plains (Philben et al., 2020).

Soil redox conditions, along with other edaphic factors such as pH and nutrient availability, are linked to microbial community composition and metabolic diversity (Philippot et al., 2023). Microbial characteristics can be assessed using metagenomics, which has proven particularly valuable in extreme environments such as thawing permafrost, where genomic analyses have revealed microbial adaptations to redox-stratified conditions and geochemical gradients (Romanowicz et al., 2023; Waldrop et al., 2023; Woodcroft et al., 2018). Romanowicz et al., 2023, for instance, showed that imicrobial iron reduction strongly influences microbial carbon degradation in thawing permafrost. Similarly, Waldrop et al., 2023 demonstrated that permafrost microbial communities and functional genes are structured by latitudinal gradients and soil geochemistry. In alpine plains, fluctuating water tables and variable oxygen conditions likely necessitate a wide microbial metabolic repertoire, enabling microorganisms to adapt their respiration strategies to the availability of TEAs (Fu et al., 2023; Keiluweit et al., 2016; Ruan et al., 2024). Alpine systems therefore express similar redox variability and potentially microbial adaptation strategies as thawing permafrost, yet integrated metagenomic assessments remain rare in alpine environments.

Here, we investigate the relationships between SOC stocks, microbial community structure and functional potential, and environmental conditions across plain and slope areas in two alpine headwater catchments. Although environmental conditions differ slightly between the two catchments, both share similar geomorphic structures and hydrological regimes and can therefore be treated as landscape-level replicates. We hypothesise that:

1. Plain soils exhibit anoxic conditions that are associated with higher SOC contents and higher levels of poorly degradable SOC, such as phenols and aromatics.
2. Microbial communities exhibit greater metabolic diversity in plain than slope soils due to larger temporal variability in soil redox conditions.

To test our hypotheses, we combine the analysis of soil physicochemical characteristics with analyses of soil redox state by mediated electrochemistry, soil organic matter chemistry by pyrolysis gas chromatography-mass spectrometry, and microbial functional diversity and metabolic capabilities by shotgun metagenomics. We compared SOC content and composition across landscape positions and soil depths, correlated taxonomic lineages with functional gene potentials, and incorporated environmental vectors into a non-metric multidimensional scaling (NMDS) ordination analysis to assess the relationships between microbial communities and environmental factors.

## Materials and Methods

### Site Description and Sample Collection

Soils were collected from the riparean areas of two natural headwater catchments in the Swiss Alps: Blatt in the Binntal valley (46°22’N/8°16’E) and Ar du Tsan (46°12’N/7°30’E) in the Vallon de Réchy. Both catchments feature a mixed bedrock mainly comprised of gneiss and carbonated rock, and diverse vegetation made up of siliceous alpine grasslands and moorlands (Frélechoux and Gallandat, 1995; Matteodo et al., 2018; Richard et al., 1993; swisstopo, 2024). The sampling sites within the two catchments were strategically chosen to encompass both slope and plain areas. At Réchy, the elevation of sampling locations varied from 2154 to 2243 meters above sea level (m a.s.l.), while at Binntal, the range was between 1984 and 2105 m a.s.l. Soil sampling was carried out in late July 2023. Average July temperatures at l’Ar du Tsan and Binntal are 12.9 °C and 8.7 °C, with precipitation levels of 76 mm and 97 mm, respectively (Fick and Hijmans, 2017). At 13 sampling locations (Figure 1), soil from three soil depths (0-10 cm, 10-30 cm, and 30-50 cm, when available - see Table S1) was collected with an auger. Precise coordinates, slope, and specific landscape position were recorded for each location. Once gathered, the samples were placed in zip-lock bags; bags for water-logged soils contained oxygen scrubbers. Samples were kept cool with ice packs, and transported to the laboratory. Samples for soil physicochemical characterization were immediately processed; sub-samples for soil redox characterization were stored at -20 °C until analysis. Samples for DNA extraction were placed in a sterile manner into Whirl-Pack® bags and homogenised directly by kneading. Post-homogenisation, the soil was split into triplicate subsamples, put into cryotubes snd shock-frozen using liquid nitrogen. After transport to the laboratory, they were stored at -80 °C until analysis.

**Figure 1:**
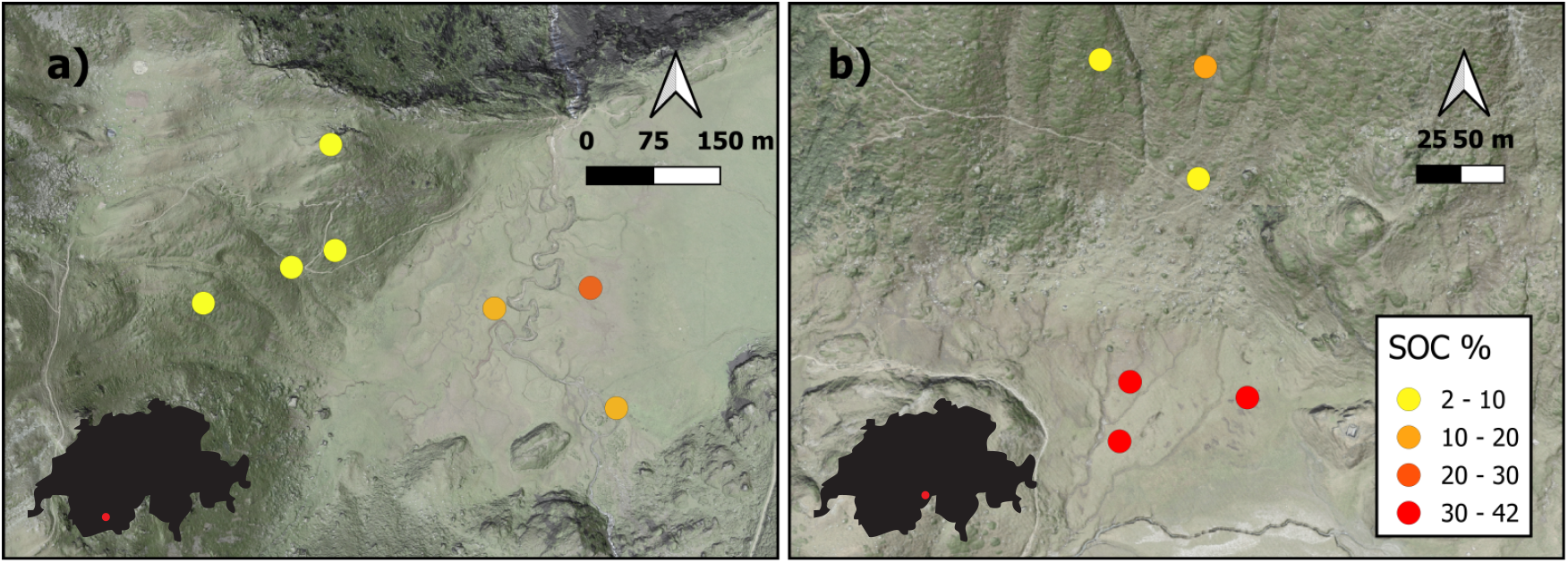
Spatial distribution of mean soil organic carbon (SOC) stocks, calculated from three soil depth-specific samples within the top 50 cm of soil, at **a** Réchy and **b** Binntal. Slope soils are represented in shaded terrain, plain soils in unshaded areas. Background: SWISSIMAGE orthophoto and hillshade from swissALTI3D.

### Soil Physicochemical Characterisation

#### Sample preparation, pH measurements, and soil texture analyses

Soil samples were air-dried and subsequently oven-dried at 105°C, homogenised, sieved through a 2 mm mesh to remove coarse particles (e.g., plant roots and stones), and ground using a ball mill (Pulverisette 7, Fritsch) to achieve a fine, uniform powder. Dry weight was determined from changes in soil mass upon drying. Soil pH was measured SevenDirect SD50 pH meter, Mettler Toledo) in a 1:5 soil-to-deionised water suspension after 30 minutes of agitation at 200 RPM, followed by 30 minutes of settling (Table S2). Soil texture was analysed using laser diffraction (LS 13 320, Beckman Coulter) with a grain size analyser, on 0.5 g of air-dried, sieved bulk soil following organic matter digestion with hydrogen peroxide over two weeks (Table S2).

#### Elemental composition

Total carbon was measured by chromatography after combustion at 900 °C on a CHNS element analyser (Flash EA 1112, Thermo Finnigan). As the soils lacked carbonates, the total SOC content (expressed as weight % of dry soil) was considered equivalent to the measured total carbon. Total content of Fe, Mn, and S were determined on 5 g of dried, sieved, and powdered soil using X-ray fluorescence spectroscopy (SPECTRO XEPOS).

#### SOC composition

The relative abundance of major compound classes were determined using pyrolysis gas chromatography–mass spectrometry (Py-GC-MS, Tolu et al., 2015). Soil samples were placed in clean, fire-polished quartz tubes and pyrolysed at 600°C for 20 seconds under a helium flow. The released pyrolysis moieties were transferred via a heated transfer line into an Agilent 7980A GC equipped with a Zebron ZB-5MS column (Phenomenex, Woerden, the Netherlands; 30 m × 250 *µ*m × 0.25 *µ*m) coupled to an Agilent 5975C MSD single quadrupole mass spectrometer operating in electron ionisation mode (scanning *m/z* 50 to 650 at 2.7 scans per second; ionisation energy: 70 eV), using helium as the carrier gas and introduced in split mode (70: 1 split ratio; constant flow of 2 ml per min, with gas saver mode active). The pyrolysis transfer line and rotor oven temperature were maintained at 325°C, the heated GC interface at 280°C, the electron ionisation source at 230°C, and the quadrupole at 150°C. The GC oven was programmed from 40°C (held for 5 minutes) to 300°C at 5°C per minute, where it was held for 3 minutes, giving a total run time of 60 minutes. Approximately 106 of the most abundant pyrolysis moieties were identified, identified by comparing their retention times and spectra to entries in the NIST Mass Spectral Library and grouped into categories based on their origin and chemical characteristics: lipids, lignins, polysaccharides, phenols, nitrogen containing compounds, and aromatics (Figure S1 and Table S1). Given the complexity of the pyrograms, it was not possible to integrate individual moiety in total ion current mode due to significant overlap between ion peaks. Instead, single ion filtering was used to measure the peak area of each compound. The major ions of each compound were filtered and integrated (Table S3). The relative abundance of each identified compound was calculated as a percentage of the total identified compounds.

#### Electron accepting & donating capacities

Electron accepting and electron donating capacities (EAC & EDC), were determined through mediated electrochemical analyses using an 8-channel potentiostat (CH Instruments, Inc.) in an anoxic environment inside a glovebox workstation (Labmaster pro MBraun), as previously described (Aeppli et al., 2018, 2022). Experiments were conducted using a pH-buffered solution at pH 5.5 (0.4 M sodium acetate-acetic acid) with 10 mM sodium chloride as a background electrolyte. Mediated electrochemical reduction (MER) potentials were set versus standard hydrogen electrodes at -0.51 V vs. SHE (*E*_H, MER_) and mediated electrochemical oxidation (MEO) potentials at +0.82 V vs. SHE (*E*_H, MEO_). For EAC measurements, ethyl viologen (Michaelis and Hill, 1933) was used as an electron transfer mediator; for EDC, ABTS (Thomas et al., 2004) was used. To prepare the samples, 1 g of frozen soil was transferred into 10 mL of Milli-Q water under anaerobic conditions to create a slurry. For each measurement, 30 µL of the slurry was used for EAC determination, and 20 µL for EDC determination. An additional 1 mL aliquot was taken from each slurry in triplicate to determine soil dry weight for normalisation. During EAC determination, electrons are delivered from the electrode via the mediator to the sample’s redox reactive constituents, thereby reducing them. Conversely, during EDC measurement, electrons are withdrawn from these constituents through the mediator and transferred to the electrode. EAC and EDC values were determined from current responses measured upon the addition of the sample to the electrochemical cells at *E*_H, MER_ and *E*_H, MEO_, respectively. Capacities (mol *e*^−^ g^−1^ dried soil) were determined by integrating the baseline-corrected current, *i*(*t*), over time from *t*_0_ until the current returned to baseline at *t*_end_ following equations 1 and 2.

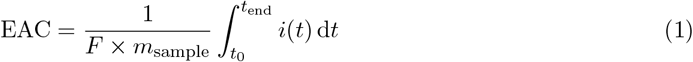

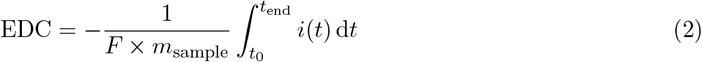

where *F* ≈ 96485 C mol^−1^ is the Faraday constant and *m*_sample_ is the dry mass of the soil.

### DNA Extraction, Sequencing, and Analysis

DNA was extracted from 0.5 g of soil using the DNeasy PowerSoil Pro Kit (Qiagen®, Germany) following the manufacturer’s protocol. DNA content and purity were assessed using microspectrophotometry (NanoDrop One; Thermo Fisher Scientific Inc., USA). Library preparation and shotgun metagenomic sequencing were performed by Novogene (UK) with Illumina NovaSeq 6000 platform to generate paired-end (150 bp) reads.

Initial quality checks of raw sequencing reads were conducted using FastQC to ensure data integrity (Andrews, 2010). Reads were subjected to quality filtering using fastp (Chen et al., 2018), followed by a second round of quality checks with FastQC to verify improvements in read quality. De novo assembly of high-quality reads was performed with MEGAHIT, generating contigs suitable for downstream analyses (Li et al., 2015). Assembly statistics were evaluated using QUAST to ensure completeness and accuracy (Gurevich et al., 2013). High-quality reads were mapped to assembled contigs using Strobealign to generate coverage profiles (Sahlin, 2022). Metagenome-assembled genomes (MAGs) were reconstructed using MetaBAT2 (Kang et al., 2019), and bin quality was assessed using CheckM2 to ensure completeness and contamination metrics were within acceptable thresholds (Chklovski et al., 2023). Taxonomic classification of MAGs was assigned using the Genome Taxonomy Database Toolkit (GTDB-Tk, Chaumeil et al., 2020). Functional annotation of MAGs was conducted using METABOLIC (Zhou et al., 2021), allowing for the prediction of key metabolic pathways and biogeochemical functions. Dereplication of MAGs was performed using dRep to consolidate redundant genomes and generate a representative set (Olm et al., 2017). The relative abundance of dereplicated MAGs was calculated using CoverM (Wood and Salzberg, 2024).

### Statistical Analysis

SOC content and composition were analysed using Wilcoxon rank-sum tests for comparison of plain and slope soils and one-way ANOVA with Tukey’s HSD post-hoc tests for soil depth effects. Spearman correlations were used to examine associations between functional gene categories and taxonomic lineages identified in the metagenomic data. Non-metric multidimensional scaling (NMDS) analysis was used to to visualise microbial community dissimilarities based on Bray-Curtis distances and identify effects of environmental factors on microbial community composition. This analysis was complemented by a PERMANOVA test to assess the influence of location, landscape position, and soil depth on microbial community structures. Figures and statistical analyses were generated in R using the `vegan` package to explore microbial community composition and diversity metrics (Oksanen et al., 2015).

## Results

### Plain Soils Store More Organic Carbon Than Slope Soils

Plain soils had higher SOC contents than slope soils at all soil depths (Figure 2): SOC ranged was 24.3 *±* 8.3% in the 0 - 10 cm layer and 22.8 ± 20.5% at 30 - 50 cm. with highest SOC values observed in the mid-soil depth layer (10 - 30 cm: 29.6 ± 16.3%) for plain soils. In contrast, slope soils exhibited a clear soil depth-dependent trend in SOC concentrations, with SOC decreasing from 6.49 ± 4.92% in the surface layer (0 - 10 cm) to 2.48 ± 0.66% at 10 - 30 cm, and 1.80 ± 0.20% at 30 - 50 cm. SOC content in plain soils was significantly higher than that of the slope across all soil depths, with mean SOC at 0 - 10 cm soil depth in plain soils being approximately 3.7 times higher than in slope soils. The spatial distribution of SOC stocks is shown in Figure 1. In both the Réchy and Binntal catchments, plain soils in unshaded terrain showed higher SOC (Réchy: 10 - 30%, Binntal: 20 - 42%), while shaded slope soils were lower (2 - 10% for both).

**Figure 2:**
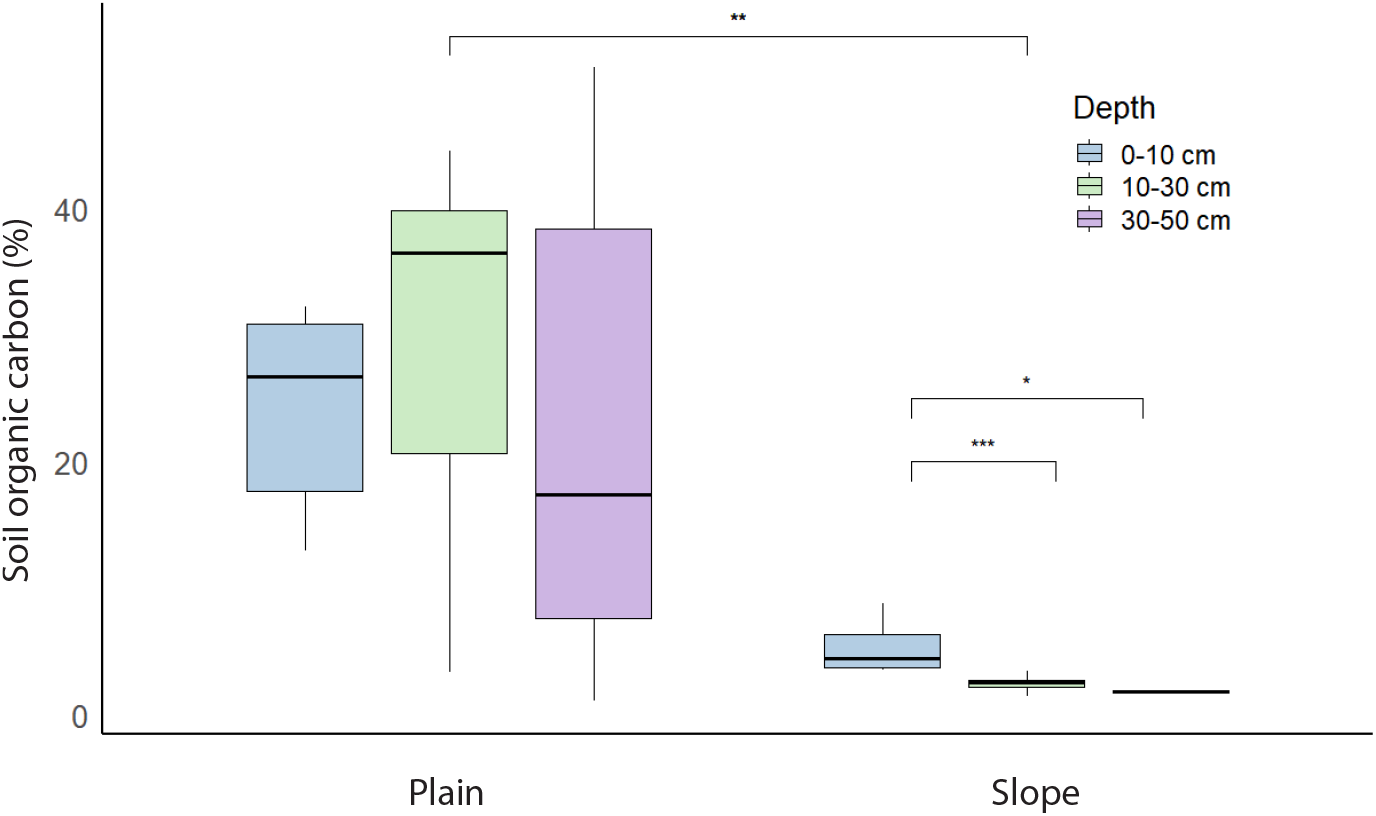
Soil organic carbon content (percentage of soil dry weight) at three soil depths in plain and slope soils (*n* = 20). Asterisks indicate levels of significance: * for *p* < 0.05, ** for *p* < 0.01, and *** for *p* < 0.001.

### Soil Organic Carbon Composition Differs by Landscape Position and Soil Depth

The composition of organic matter varied by landscape position and soil depth (Figure 3). Slope soils were enriched in polysaccharides (31.9 %), whereas plain soils contained higher proportions of phenols (20.9 %). The proportions of aromatics, lipids, nitrogen-containing compounds, and lignin was comparable between both types of soils. The relative contribution of aromatic compounds increased with soil depth from 23.3 % at 0 - 10 cm to 36.4 % at 30 - 50 cm, while the relative contriubtion of lignins decreased from 8.2 % at 0 - 10 cm to 2.1 % at 30 - 50 cm.

**Figure 3:**
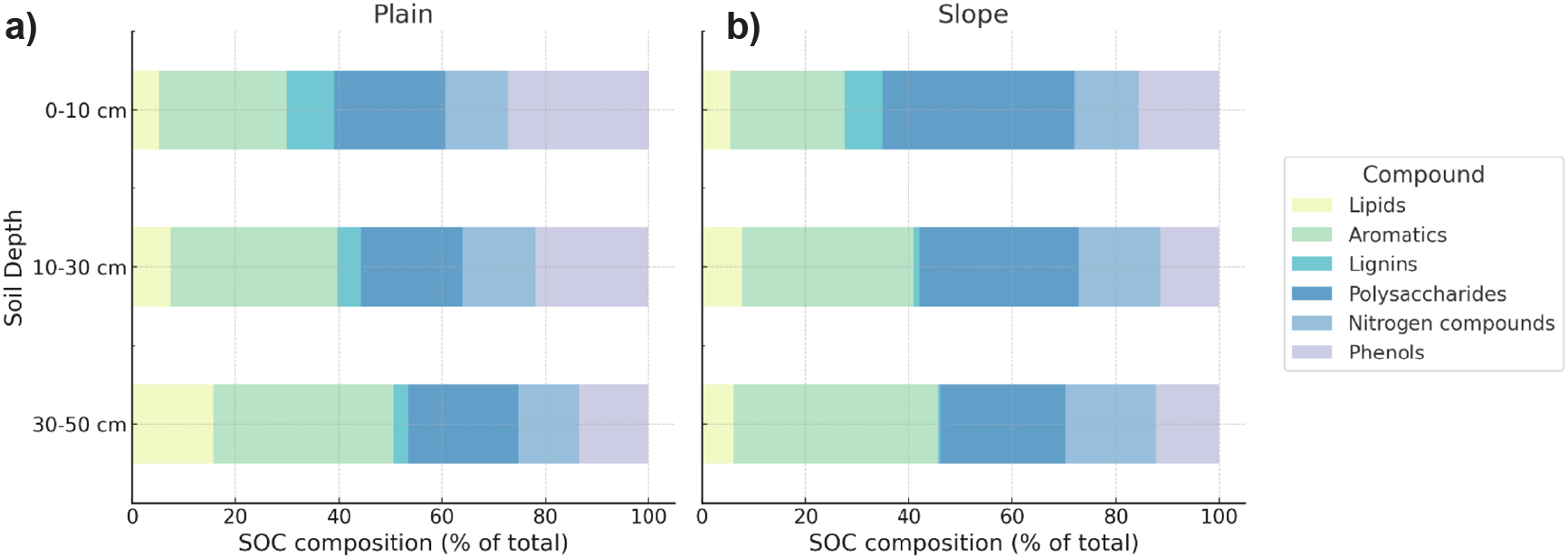
Composition of soil organic carbon (SOC) in plain (**a**) and slope soils (**b**). Relative abundances for each compound class are provided in Table S1.

### Plain Soils Exhibit soil depth-Dependent Redox Zonation

Soil redox state was described the EAC and EDC values, which represent the contribution of the soils’ pools of redox-active oxidised and reduced geochemical species, respectively. In plain soils, EAC and EDC values exhibited an inverse relationship, with EDC increasing from 0.12 ± 0.05 to 0.33 ± 0.07 mmol g^−1^ soil and EAC decreasing from 0.41 ± 0.10 to 0.16 ± 0.05 mmol g^−1^ soil from 0-10 cm to 30-50 cm soil depth, consistent with increasingly reducing conditions (Figure 4a). In contrast, slope soils showed low EDC values (e.g., 0.02 ± 0.01 at 10 cm) and constant EAC values (e.g., 0.48 ± 0.12 at 10 cm) across all soil depths, indicating oxic conditions (Figure 4b). We compared total electron exchanging capacity (sum of EAC and EDC) to elemental composition to attribute the EAC and EDC responses to geochemical phases (Figure S2). In plain soils, iron explained most of the redox reactivity, followed by sulfur. In slope soils, iron was the dominant redox-active phase, with a small contribution from redox-active organic matter.

**Figure 4:**
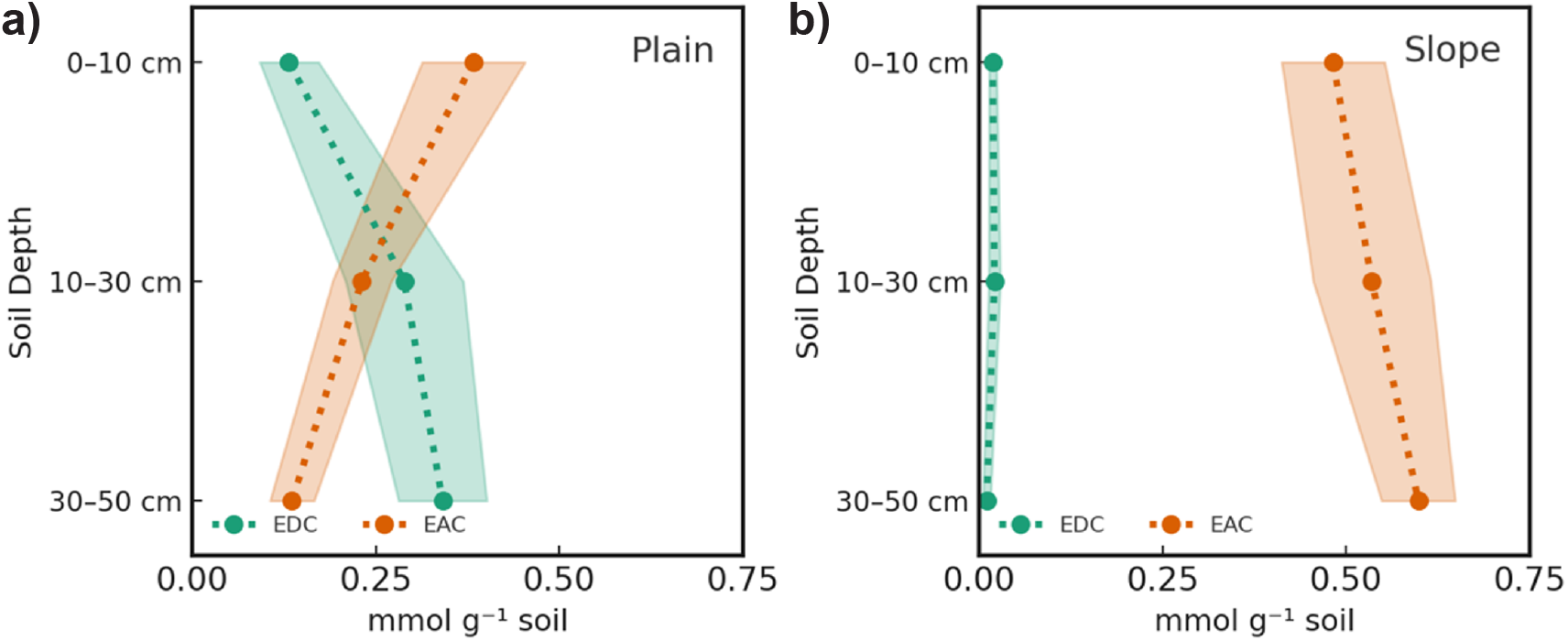
Average electron donating (EDC, green) and accepting capacities (EAC, orange) of plain (**a**) and slope soils (**b**). Shaded areas represent the standard error of the mean. Each data point reflects the average of five soil samples.

### Microbial Community Composition and Functional Potential Are Linked to Soil Redox Conditions

Microbial community composition differed between plain and slope soils (Figure 5): plain soils had higher relative abundances of *Chloroflexota, Acidobacteriota*, and *Desulfobacterota*, whereas slope soils contained greater proportions of *Verrucomicrobiota, Thermoproteota*, and *Dormibacterota*. The heat map of functional genes (Figure 6) shows that plain soils exhibited higher relative abundances of genes assigned to nitrate, metal (Fe/Mn), and sulfate reduction across all soil depths, while slope soils had lower relative abundances of these genes. Potential methanogenesis genes were not detected, whereas genes attributed to methane oxidation (*Methylomirabilota*) were present in both landscape positions at low abundance.

**Figure 5:**
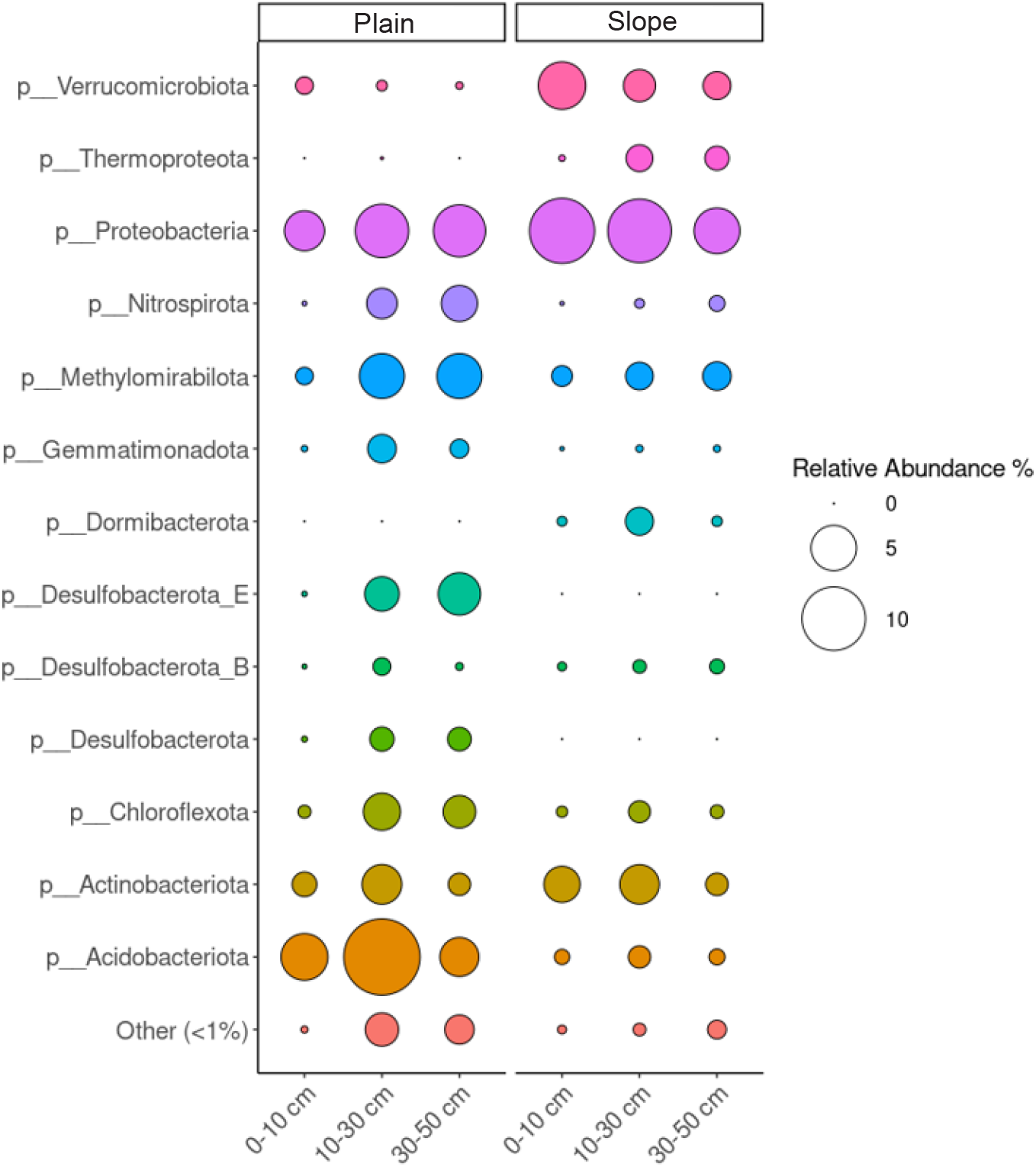
Microbial community composition. The relative abundance of key prokaryotic phyla is shown for plain and slope soils at three soil depths. Circle size represents the relative abundance of each taxon.

**Figure 6:**
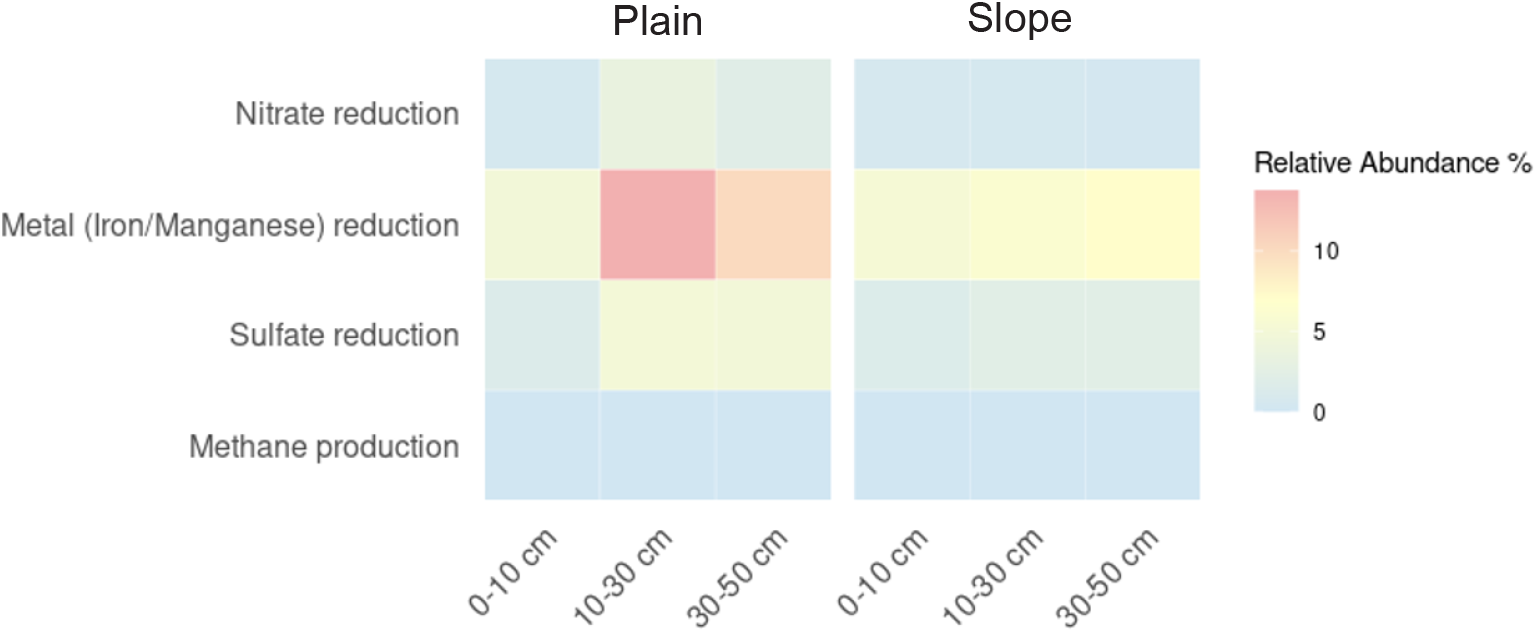
Heatmap of potential anaerobic microbial respiration pathways in plain and slope soils across three soil depths. Genes associated with nitrate, iron/manganese, and sulfate reduction, as well as methane production, are shown as relative abundances.

Taxon–function links were identified between specific microbial lineages and key reductive processes (Table 1). *Chloroflexota* lineages (e.g., class *Anaerolineae*; class *Dehalococcoidia* lineages DSTF029 and SM23-31) correlated with nitrate-reduction genes. *Acidobacteriota* classes *Thermoanaerobaculia, Acidobacteriae*, and *Blastocatellia* correlated with Fe/Mn-reduction genes. Orders within *Desulfobacterota* (*Geobacterales*, BSN033, and *Desulfatiales* ) correlated with sulfate-reduction genes.

**Table 1:**
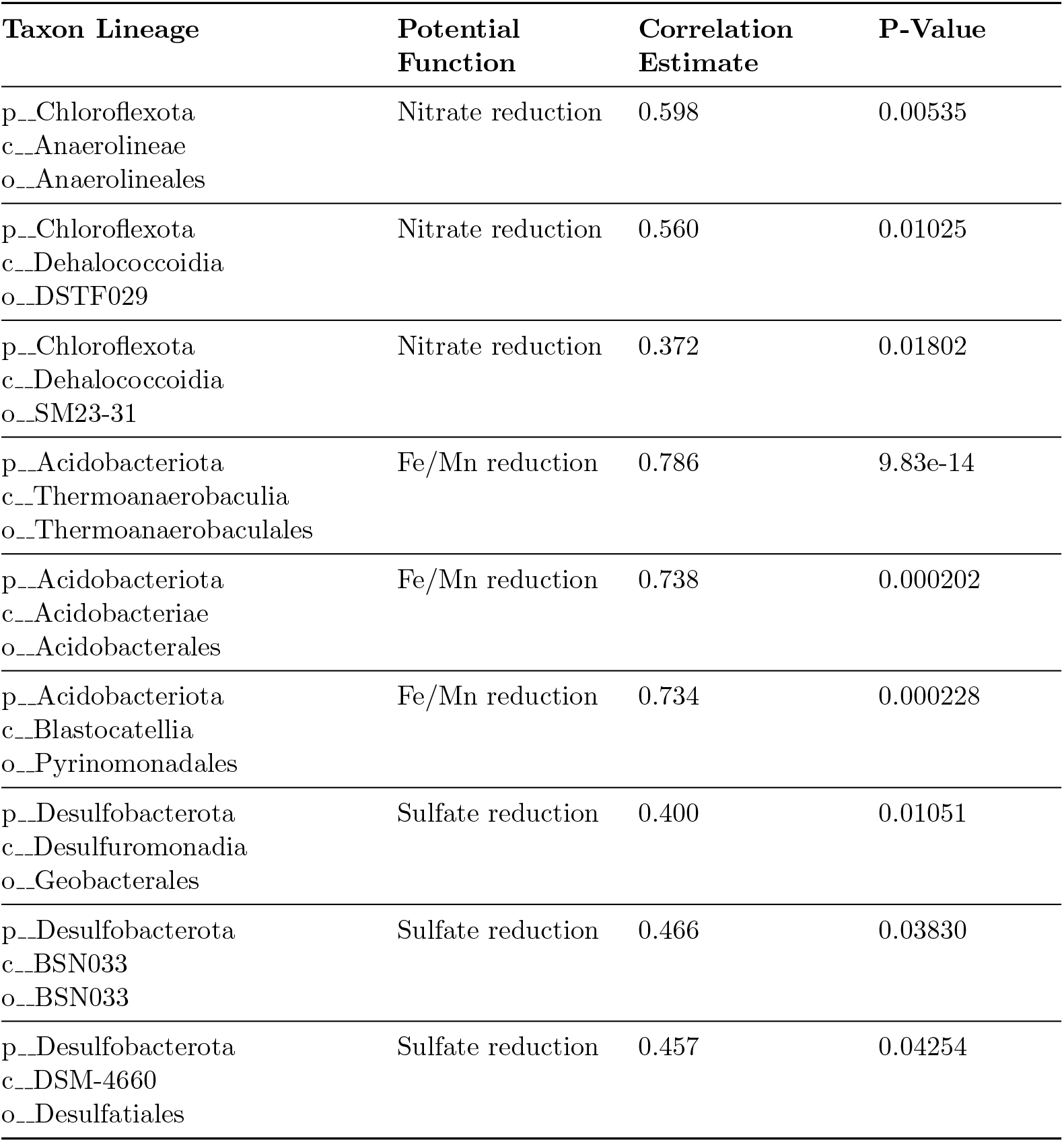
Spearman correlations between functional gene categories and taxonomic lineages identified in metagenomic data. Lineages are grouped by taxonomic rank: p = phylum, c = class, o = order. Only statistically significant associations (*p* < 0.05) are shown.

### Microbiomes Exhibit Enhanced Metabolic Versatility and Greater Potential for Anaerobic Carbon Turnover in Plain Soils

We assessed the core set of metabolic functions relative to carbon turnover following the flowgram pipeline by Zhou et al., 2021. Distribution of these functions, showing the proportion of metagenome-assembled genomes (MAGs) that encode the genes required for each transformation, indicate that SOC oxidation genes were present in 24.67 % of plain soil genomes (444 MAGs) versus 18.94 % in slope soil genomes (Figure 7). Fermentation potential was likewise greater in plain soil communities (12.85 %; 234 MAGs) than in slope soil communities (8.10 %). Hydrogen generation genes occurred in 11.61 % of plain soil genomes (231 MAGs) compared with 7.51 % of slope soil genomes, whereas hydrogen oxidation genes were found in 4.47 % and 1.30 % of genomes, respectively (74 MAGs). In contrast, acetate-oxidation genes showed slightly higher representation in slope soil communities (1.78 %; 28 MAGs) than in plain soil communities (1.30 %) while genes for ethanol oxidation and carbon fixation were detected at low levels in both settings (ethanol oxidation: 10.35 % plain, 9.55 % slope; carbon fixation: 1.09 % plain, 0.40 % slope). Methanotrophy genes were rare but detectable (0.66 % plain, 0.76 % slope; 21 MAGs), whereas methanogenesis genes were absent from all assemblies. Overall, the higher prevalence of fermentation, hydrogen metabolism, and SOC-oxidation genes in plain soil MAGs indicates a larger genomic investment in anaerobic carbon turnover than in slope soils.

**Figure 7:**
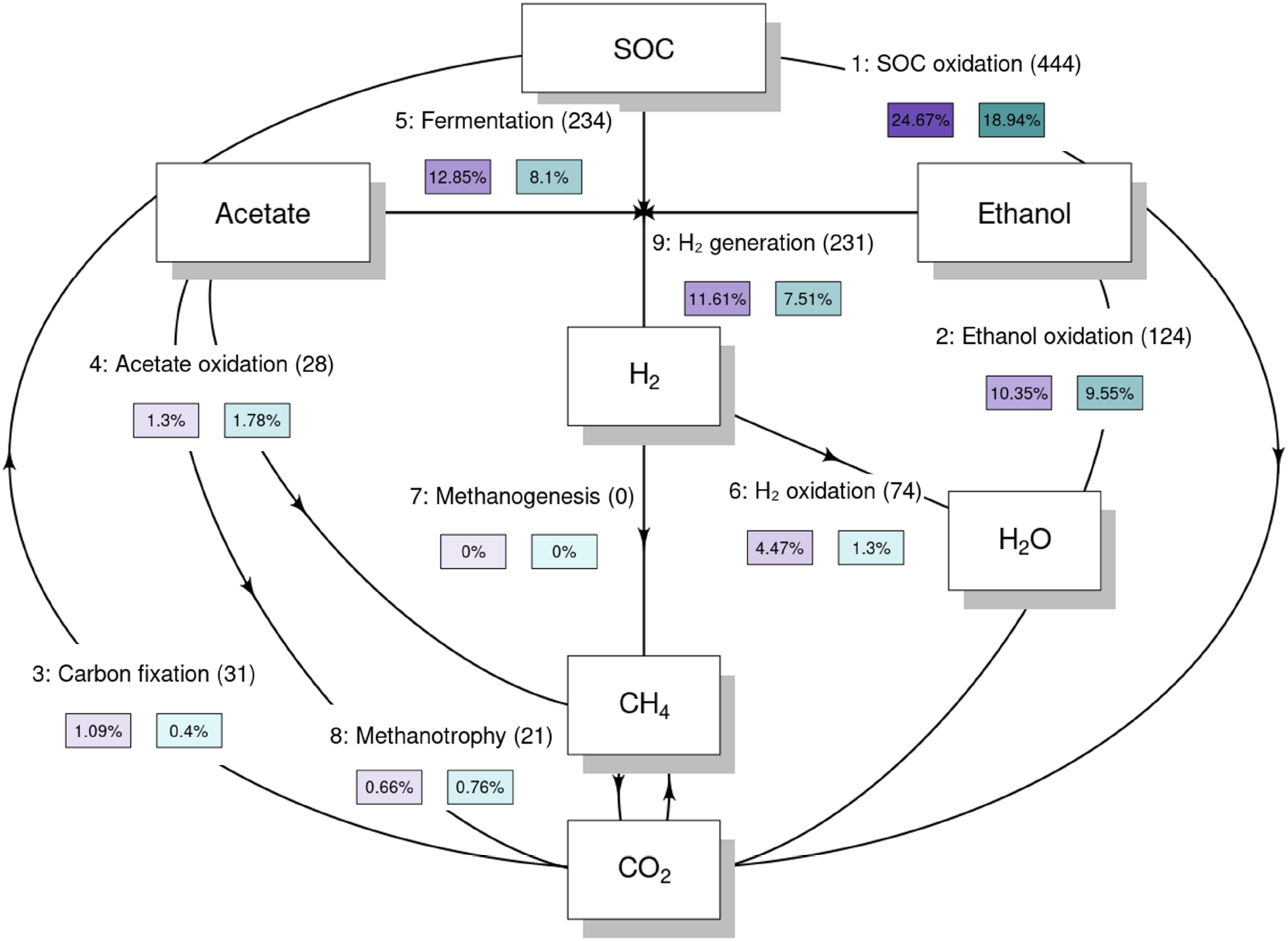
Soil organic carbon (SOC) transformations mediated by microbial communities in plain and slope soils. The flowgram illustrates SOC-related metabolic steps reconstructed from metagenomic data using a modified script from METABOLIC (Zhou et al., 2021). Each arrow represents a distinct transformation step, with boxes denoting key compounds involved. Arrow labels indicate the step number and transformation type, the number of genomes encoding the necessary genes (in brackets), and the relative abundance of those genomes in plain (purple) and slope soil communities (teal), expressed as a percentage of total community composition. Community-level genome abundance and function were inferred from metagenome-assembled genomes.

### Microbial Community Composition Correlates with Soil Physicochemical Properties Across Landscape Positions and Catchments

Similarities of microbial community composition across catchments and landscape positions were assessed using an NMDS plot (Figure 8, supplementary environmental values in Table S2). Plain soil communities cluster on the bottom left, while slope soil communities cluster on the top right with minimal overlap between groups. Vectors for Ca content, soil pH, EAC, and phenols have the largest percentage value and point toward the plain soil cluster, indicating strong positive correlations with those communities. Conversely, vectors for polysaccharides, nitrogen-containing compounds, EDC, and clay content project toward the slope soil cluster. Sand and silt vectors plot between the two groups with intermediate vector lengths. Thus, variation in Ca content, pH, EAC, EDC, and specific SOC fractions (phenols, nitrogen compounds, and polysaccharides) aligns with the primary ordination axis that separates plain and slope microbial assemblages.

**Figure 8:**
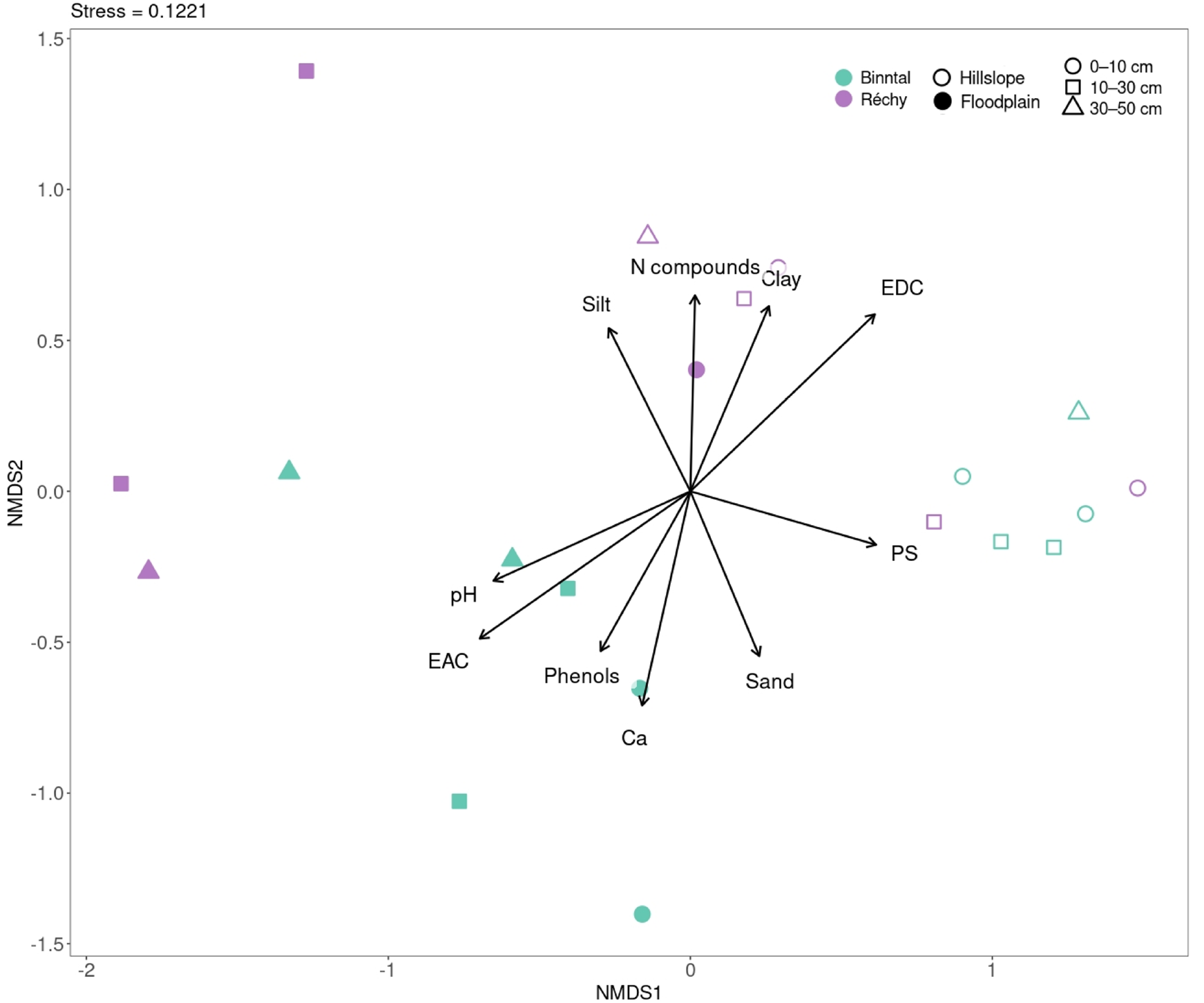
NMDS plot illustrating the microbial community composition of soils from two alpine headwater catchments, Binntal and Réchy, based on Bray-Curtis dissimilarity. Environmental vectors overlaid on the ordination indicate the direction and strength of correlations between environmental variables and microbial community composition. Vector length is scaled by the square root of the r^2^ value, reflecting the strength of these correlations. The vectors represent the top ten environmental variables, selected based on descending r^2^ values from envfit analysis. These variables include polysaccharides (PS), nitrogen compounds (N compounds), silt, clay, sand, soil pH, electron accepting capacity (EAC), calcium (Ca), phenols, and electron donating capacity (EDC). PERMANOVA attributes 23.3 % of the Bray–Curtis variation to landscape position (*R*^2^ = 0.2325, *p* = 0.001) and 10.0 % to catchment identity (*R*^2^ = 0.0999, *p* = 0.030); soil depth effects are negligible.

## Discussion

### Soil Redox State Is Linked to Soil Organic Carbon Quantity and Chemistry

Plain soils had substantially higher SOC content than slope soils, with nearly four times more carbon per g dry soil in the surface layer (0 - 10 cm; Figure 2). These differences mirrored differences in soil redox state (Figure 4). In plain soils, conditions became increasingly reducing with soil depth. The observed EDC values likely reflect the accumulation of reduced organic compounds, ferrous iron, and sulfide, produced vial microbial respiration under past anoxic conditions. Iron was the major contributor to electron exchanging capacity and was therefore a key TEA in plain soils. In slope soils, no EDC was detected, suggesting that these soils were fully oxic. Most of the EAC response was explained by iron with some contribution from organic matter. The observed patterns in SOC content and soil redox state are therefore in agreement with our first hypothesis stating that soils on the plain exhibit anoxic conditions that are associated with higher SOC contents.

Differences in SOC composition between plain and slope soils are likely due to variations in organic matter inputs and preservation mechanisms. Plain soils were enriched in phenolic compounds, whereas slope soils contained higher proportions of polysaccharides (Figure 3). This pattern was linked to contrasting vegetation types and moisture regimes. On the plain, grasses and sedges produce litter rich in phenol-containing structural polymers, which are selectively preserved under periodically anoxic conditions because the degradation of phenolic compounds depends on extracellular oxidative enzymes, such as phenol oxidase and peroxidase (Fenner and Freeman, 2011; Freeman et al., 2001). In contrast, slope soils dominated by dwarf shrubs and upland herbs receive litter rich in easily degradable carbohydrates, which likely causes the higher abundance of polysaccharides. Across both landscape positions, we observed a decline in lignin-derived compounds and an increase in aromatic compound contributions with soil depth. The relative higher lignin content in surface soils likely reflects recent plant inputs from vascular tissue. Aromatic compounds are chemically more stable and persist under oxygen-limited conditions (Fenner and Freeman, 2011; Freeman et al., 2001; Sinsabaugh, 2010). Combined, these findings indicate that as soil depth increases, lignin is progressively broken down or transformed, while less bioavailable aromatic structures accumulate.

The observed trends in SOC content and composition, and soil redox state align with the expected sequential microbial use of substrates based on reaction thermodynamics. Compared to polysaccharides, phenolic and aromatic compounds are chemically more reduced on average and therefore require higher energy input to be oxidized. Under anoxic conditions, this required energy input may outweigh the energy released upon redeuction of alternative TEAs, resulting in the accumulation of these compounds (Boye et al., 2017; Gunina and Kuzyakov, 2022).

### Microbial Community Composition and Potential Functions are Linked to Landscape Position and Soil Physicochemical Characteristics

Microbial community composition differed between plain and slope soils (Figure 5), with plain soils having higher relative abundances of potential microbial respiratory pathways, in agreement with our second hypothesis. These higher abundances are likely driven by seasonal moisture fluctuations and nutrient-rich conditions. Recurrent anoxic windows likely support a rich assemblage of anaerobic microbial metabolisms, including the potential reduction of nitrate, iron, manganese and sulfate, thereby sustaining SOC turnover in the riparian corridor (Keiluweit et al., 2016). We found higher abundance of taxa commonly associated with anaerobic respiration at soil depth in plain soils, including members of the *Chloroflexota* or *Desulfobacterota* phyla (Eilers et al., 2012), in line with the redox stratification inferred from EDC–EAC profiles. Conversely, slope soils were dominated by phyla such as *Verrucomicrobiota, Thermoproteota* and *Dormibacterota*. These soils were well-drained and oxygen-rich which, in concert with lateral inputs of carbohydrate-rich litter, promotes high-energy yielding aerobic microbial respiration pathways. Methanogenesis genes were detected neither in plain nor slope soils; however, low-abundance methanotrophs (*anaerobic Methylomirabilota*) were present in both soils, implying that any methane produced in deeper plain soils or in anoxic microsites in slope soil was oxidised. The absence of canonical mcrA-bearing MAGs does not necessarily preclude methanogenesis; these genes are often confined to low-abundance taxa inhabiting deeper, more water-logged soil horizons and can evade detection when reads assemble into short or unbinned contigs (Woodcroft et al., 2018). Differential expression analyses further highlighted up-regulation of Group 3 and Group 1 NiFe hydrogenases in plain soils (Figure S3), enzymes that confer redox plasticity by allowing microbial communities to alternate between fermentative and respiratory metabolic strategies as oxygen availability fluctuates (Piché-Choquette and Constant, 2019).

Microbial community composition aligned with the observed differences in SOC composition across landscape positions (Figure 5). Microbial community assemblages in plain soils likely need to invest more in enzymatic machinery for the partial degradation and preservation of phenol-rich litter derived from hydrophilic grasses and sedges (Piché-Choquette and Constant, 2019; Yang et al., 2021). The selective preservation of phenolic compounds under anoxic conditions likely reflects slow decomposition of soil organic matter, and these compounds may also contribute to the elevated EDC observed in plain soils by providing redox-active moieties that function as extracellular electron shuttles (Fenner and Freeman, 2020; Kappler et al., 2004). In contrast, microbial communities in slope soils were dominated by fast-growing copiotrophs and likely preferred polysaccharide-rich substrates, congruent with the higher proportion of depolymerisable carbohydrates we observed in these soils. Building on the previous discussion, the abundance of aromatic compounds increased with soil depth, whereas lignin-derived phenols decreased. Surface soil horizons may therefore support microbial communities that favour easily oxidisable polysaccharides and secrete ligninolytic oxidases, driving rapid lignin turnover (Baldrian, 2017; Dao et al., 2022).

Several environmental variables influenced microbial community structure in plain and slope soils (Figure 8). EAC, Ca content, pH and phenolic content explained most variance in microbial community composition in plain soils. Calcium has previously been shown to stabilise SOC through cation bridging with negatively charged organic surfaces, potentially restricting microbial access to SOC (Rowley et al., 2018). Given the pH-dependence of these interactions and the propensity of alpine plain systems to experience seasonal water saturation (Ma et al., 2021), it is plausible that associated redox and pH fluctuations influenced microbial niche differentiation (Blagodatskaya and Kuzyakov, 2008). The observed association between phenolic content and microbial community composition may also reflect the presence of redox-active substrates that select for microbial groups capable of utilising these substrates either as carbon sources or as electron shuttles under oxygen-limited conditions. In slope soils, microbial assemblages were more closely aligned with polysaccharide content, EDC, and clay content. While mean EDC values were relatively low, the spatial variation across samples may point to micro-heterogeneity in the distribution of redox-active substrates, which could influence microbial organisation even under well-drained conditions. Polysaccharides, derived from rapid cycling of plant litter, may provide readily accessible energy, selecting for copiotrophic lineages (notably several proteobacterial MAGs that our metagenomic analysis showed to be enriched in slope soils). Clay content may also play a role through its known capacity to stabilise organic matter via adsorption, thereby modifying substrate accessibility (Sollins et al., 1996). Our results suggest that substrate quality, rather than the amount of SOC only, shape microbial assemblages under well-aerated conditions.

## Conclusions

Our work shows how microbial community composition varies across landscape positions from wet plain soils to drier slope soils in alpine riparian zones. Plain soils contained three to four times more SOC than adjacent slope soils, were enriched in phenolic compounds, had higher EDC values, and harbored microbial communities with genes for nitrate, iron, manganese, and sulfate reduction—features consistent with periodic anoxia and the accumulation of SOC due to thermodynamic limitations on microbial activity. In contrast, slope soils had lower SOC contents, were not reduced, had a higher proportion of labile polysaccharides, and microbial communities dominated by aerobic taxa. Together, these patterns demonstrate how moisture-driven redox regimes shape microbial potential and SOC composition, influencing the balance between SOC preservation and mineralization across the landscape. By comparing analogous landscape positions in two independent alpine catchments, our work provides a case study of how topographically driven redox gradients govern microbial ecology.

Several open questions remain regarding the role of microbial metabolism in carbon cycling in alpine riparian soils, particularly during seasonal transitions. Microbial communities may remain active beneath the winter snowpack, but their response to the spring melt pulse of dissolved organic carbon is not well understood. Future studies applying metatranscriptomics, extracellular enzyme assays, and stable isotope probing could capture these dynamics, revealing how microbial activity under shifting redox and substrate conditions influences SOC stability. Such insights would clarify how seasonal and topographic variability regulates organic carbon turnover and ultimately the net carbon balance of alpine catchments.

## Acknowledgements

The authors thank Lorenz Schwab, Antoine Wallart, and Eric Pizem for their support with soil sampling, and Gordanna Pistoletti for technical assistance. We also thank the Swiss National Science Foundation for financial support (Grant No. 212056).

## Notes

### Competing Interest Statement

The authors have declared no competing interest.

## References

Aeppli, M., Thompson, A., Dewey, C., & Fendorf, S. (2022). Redox properties of solid phase electron acceptors affect anaerobic microbial respiration under oxygen-limited conditions in floodplain soils. Environmental Science & Technology, 56 (23), 17462–17470. 10.1021/acs.est.2c05797

Aeppli, M., Voegelin, A., Gorski, C. A., Hofstetter, T. B., & Sander, M. (2018). Mediated electrochemical reduction of iron (oxyhydr-)oxides under defined thermodynamic boundary conditions. Environmental Science & Technology, 52 (2), 560–570. 10.1021/acs.est.7b04411

Andrews, S. (2010). FastQC: A quality control tool for high throughput sequence data. https://www.bioinformatics.babraham.ac.uk/projects/fastqc/

Baldrian, P. (2017). Microbial activity and the dynamics of ecosystem processes in forest soils. Current Opinion in Microbiology, 37, 128–134. 10.1016/j.mib.2017.06.008

Berhe, A. A., & Kleber, M. (2013). Erosion, deposition, and the persistence of soil organic matter: Mechanistic considerations and problems with terminology [Commentary]. Earth Surface Processes and Landforms. 10.1002/esp.3408

Blagodatskaya, E., & Kuzyakov, Y. (2008). Mechanisms of real and apparent priming effects and their dependence on soil microbial biomass and community structure: Critical review. Biology and Fertility of Soils, 45 (2), 115–131. 10.1007/s00374-008-0334-y

Boye, K., Noël, V., Tfaily, M. M., Bone, S. E., Williams, K. H., Bargar, J. R., & Fendorf, S. (2017). Thermodynamically controlled preservation of organic carbon in floodplains. Nature Geoscience, 10 (6), 415–419. 10.1038/ngeo2940

Chaumeil, P.-A., Mussig, A. J., Hugenholtz, P., & Parks, D. H. (2020). Gtdb-tk: A toolkit to classify genomes with the genome taxonomy database. Bioinformatics, 36 (6), 1925–1927. 10.1093/bioinformatics/btz848

Chen, S., Zhou, Y., Chen, Y., & Gu, J. (2018). Fastp: An ultra-fast all-in-one fastq preprocessor. Bioinformatics, 34 (17), i884–i890. 10.1093/bioinformatics/bty560

Chklovski, A., Parks, D. H., Woodcroft, B. J., & Tyson, G. W. (2023). Checkm2: A rapid, scalable and accurate tool for assessing microbial genome quality using machine learning. Nature Methods, 20, 1203–1212. 10.1038/s41592-023-01940-w

Dao, T. T., Mikutta, R., Sauheitl, L., Gentsch, N., Shibistova, O., Wild, B., Schnecker, J., Bárta, J., Čapek, P., Gittel, A., Lashchinskiy, N., & Urich, T. (2022). Lignin preservation and microbial carbohydrate metabolism in permafrost soils. JGR Biogeosciences, 127 (1), e2020JG006181. 10.1029/2020JG006181

Davidson, E. A., & Janssens, I. A. (2006). Temperature sensitivity of soil carbon decomposition and feedbacks to climate change. Nature, 440 (7081), 165–173. 10.1038/nature04514

Duan, X., Li, Z., Li, Y., Yuan, H., Gao, W., Chen, X., Ge, T., Wu, J., & Zhu, Z. (2023). Iron–organic carbon associations stimulate carbon accumulation in paddy soils by decreasing soil organic carbon priming. Soil Biology and Biochemistry, 179, 108972. 10.1016/j.soilbio.2023.108972

Eilers, K. G., Debenport, S., Anderson, S., & Fierer, N. (2012). Digging deeper to find unique microbial communities: The strong effect of depth on the structure of bacterial and archaeal communities in soil. Soil Biology and Biochemistry, 50, 58–65. 10.1016/j.soilbio.2012.03.011

Fenner, N., & Freeman, C. (2020). Woody litter protects peat carbon stocks during drought. Nature Climate Change, 10 (4), 363–369. 10.1038/s41558-020-0727-y

Fenner, N., & Freeman, C. (2011). Drought-induced carbon loss in peatlands. Nature Geoscience, 4 (12), 895–900. 10.1038/ngeo1323

Fick, S. E., & Hijmans, R. J. (2017). WorldClim 2: New 1-km spatial resolution climate surfaces for global land areas [ eprint: https://onlinelibrary.wiley.com/doi/pdf/10.1002/joc.5086]. International Journal of Climatology, 37 (12), p4302–4315. 10.1002/joc.5086

Freeman, C., Ostle, N., & Kang, H. S. (2001). An enzymic ‘latch’ on a global carbon store. Nature, 409 (6817), 149. 10.1038/35051650

Frélechoux, F., & Gallandat, J.-D. (1995). Flore et végétation du haut val de binn entre chiestafel et le col de l’albrun [Postprint]. Bulletin de la Murithienne, 113, 105–128.

Fu, L., Xie, R., Ma, D., Zhang, M., & Liu, L. (2023). Variations in soil microbial community structure and extracellular enzymatic activities along a forest–wetland ecotone in high-latitude permafrost regions. Ecology and Evolution, 13 (6), e10205. 10.1002/ece3.10205

Gunina, A., & Kuzyakov, Y. (2022). From energy to (soil organic) matter. Global Change Biology, 28 (7), 2169–2182. 10.1111/gcb.16071

Gurevich, A., Saveliev, V., Vyahhi, N., & Tesler, G. (2013). Quast: Quality assessment tool for genome assemblies. Bioinformatics, 29 (8), 1072–1075. 10.1093/bioinformatics/btt086

Kang, D. D., Li, F., Kirton, E., Thomas, A., Egan, R., An, H., & Wang, Z. (2019). Metabat 2: An adaptive binning algorithm for robust and efficient genome reconstruction from metagenome assemblies. PeerJ, 7, e7359. 10.7717/peerj.7359

Kappler, A., Benz, M., Schink, B., & Brune, A. (2004). Electron shuttling via humic acids in microbial iron(iii) reduction in a freshwater sediment. FEMS Microbiology Ecology, 47 (1), 85–92. 10.1016/S0168-6496(03)00245-9

Keiluweit, M., Nico, P. S., Kleber, M., & Fendorf, S. (2016). Are oxygen limitations under recognized regulators of organic carbon turnover in upland soils? Biogeochemistry, 127 (2), 157–171. 10.1007/s10533-015-0180-6

Körner, C. (2021). The alpine life zone. Springer International Publishing. 10.1007/978-3-030-59538-82

Li, D., Liu, C. M., Luo, R., Sadakane, K., & Lam, T. W. (2015). Megahit: An ultra-fast single-node solution for large and complex metagenomics assembly via succinct de bruijn graph. Bioinformatics, 31 (10), 1674–1676. 10.1093/bioinformatics/btv033

Ma, M., Zhu, Y., Wei, Y., & Zhao, N. (2021). Soil nutrient and vegetation diversity patterns of alpine wetlands on the qinghai-tibetan plateau. Sustainability, 13 (11), 6221. 10.3390/su13116221

Matteodo, M., Grand, S., Sebag, D., Rowley, M. C., Vittoz, P., & Verrecchia, E. P. (2018). Decoupling of topsoil and subsoil controls on organic matter dynamics in the Swiss Alps. Geoderma, 330, 41–51. 10.1016/j.geoderma.2018.05.011

Michaelis, L., & Hill, E. S. (1933). The viologen indicators. Journal of General Physiology, 16 (6), 859–873. 10.1085/jgp.16.6.859

Oksanen, J., Blanchet, F. G., Friendly, M., Kindt, R., Legendre, P., McGlinn, D., Minchin, P. R., O’Hara, R. B., Simpson, G. L., Solymos, P., Stevens, M. H. H., Szoecs, E., & Wagner, H. (2015). Vegan: Community ecology package [R package version 2. 6-4]. https://CRAN.R-project.org/package=vegan

Olm, M. R., Brown, C. T., Brooks, B., & Banfield, J. F. (2017). Drep: A tool for fast and accurate genomic comparisons that enables improved genome recovery from metagenomes through dereplication. The ISME Journal, 11 (12), 2864–2868. 10.1038/ismej.2017.126

Pacific, V. J., McGlynn, B. L., Riveros-Iregui, D., & Welsch, D. L. (2011). Landscape structure, groundwater dynamics, and soil water content influence soil respiration across riparian-hillslope transitions in the tenderfoot creek experimental forest, montana. Hydrological Processes, 25 (5), 811–827. 10.1002/hyp.7870

Philben, M., Taş, N., Chen, H., Wullschleger, S. D., Kholodov, A., Graham, D. E., & Gu, B. (2020). Influences of hillslope biogeochemistry on anaerobic soil organic matter decomposition in a tundra watershed [First published: 29 April 2020]. Journal of Geophysical Research: Biogeosciences, 125 (7), e2019JG005512. 10.1029/2019JG005512

Philippot, L., Chenu, C., Kappler, A., Rillig, M. C., & Fierer, N. (2023). The interplay between microbial communities and soil properties [Publisher: Nature Publishing Group]. Nature Reviews Microbiology, 1–14. 10.1038/s41579-023-00980-5

Piché-Choquette, S., & Constant, P. (2019). Molecular hydrogen, a neglected key driver of soil biogeochemical processes. Applied and Environmental Microbiology, 85 (7), e02418–18. 10.1128/AEM.02418-18

Richard, J.-L., Bressoud, B., Buttler, A., Duckert, O., & Gallandat, J.-D. (1993). Carte de la végétation de la région val de réchy-sasseneire (objet cpn 3.77, alpes valaisannes, suisse) [Postprint]. Bulletin de la Murithienne, 111, 9–40.

Romanowicz, K. J., Crump, B. C., & Kling, G. W. (2023). Genomic evidence that microbial carbon degradation is dominated by iron redox metabolism in thawing permafrost [Number: 1 Publisher: Nature Publishing Group]. ISME Communications, 3 (1), 1–11. 10.1038/s43705-023-00326-5

Rowley, M. C., Grand, S., & Verrecchia, É. P. (2018). Calcium-mediated stabilisation of soil organic carbon. Biogeochemistry, 137 (1), 27–49. 10.1007/s10533-017-0410-1

Ruan, Y., Ling, N., Jiang, S., Jing, X., He, J.-S., Shen, Q., & Nan, Z. (2024). Warming and altered precipitation independently and interactively suppress alpine soil microbial growth in a decadal-long experiment. eLife, 12, RP89392. 10.7554/eLife.89392

Sahlin, K. (2022). Strobealign: Flexible seed size enables ultra-fast and accurate read alignment. Genome Biology, 23 (1), 260. 10.1186/s13059-022-02831-7

Schimel, J. P., & Schaeffer, S. M. (2012). Microbial control over carbon cycling in soil. Frontiers in Microbiology, 3, 348. 10.3389/fmicb.2012.00348

Sinsabaugh, R. L. (2010). Phenol oxidase, peroxidase and organic matter dynamics of soil. Soil Biology and Biochemistry, 42 (3), 391–404. 10.1016/j.soilbio.2009.10.014

Sollins, P., Homann, P., & Caldwell, B. A. (1996). Stabilization and destabilization of soil organic matter: Mechanisms and controls. Geoderma, 74 (1-2), 65–105. 10.1016/S0016-7061(96)00036-5

swisstopo. (2024). Swiss federal office of topography: Topographic and geospatial data [© swisstopo (2024)]. https://www.swisstopo.admin.ch

Thomas, J. H., Drake, J. M., Paddock, J. R., & Conklin, S. D. (2004). Characterization of abts at a polymer-modified electrode. Electroanalysis, 16 (7), 547–555. 10.1002/elan.200302862

Tolu, J., Gerber, L., Boily, J.-F., & Bindler, R. (2015). High-throughput characterization of sediment organic matter by pyrolysis–gas chromatography/mass spectrometry and multivariate curve resolution: A promising analytical tool in (paleo)limnology. Analytica Chimica Acta, 880, 93–102. 10.1016/j.aca.2015.03.043

Waldrop, M. P., Chabot, C. L., Liebner, S., Holm, S., Snyder, M. W., Dillon, M., Dudgeon, S. R., Douglas, T. A., Leewis, M.-C., Walter Anthony, K. M., McFarland, J. W., Arp, C. D., Bondurant, A. C., Taş, N., & Mackelprang, R. (2023). Permafrost microbial communities and functional genes are structured by latitudinal and soil geochemical gradients [Publisher: Nature Publishing Group]. The ISME Journal, 17 (8), 1224–1235. 10.1038/s41396-023-01429-6

Wood, D. E., & Salzberg, S. L. (2024). Coverm: Read alignment statistics for metagenomics. Bioinformatics. 10.1093/bioinformatics/btaf147

Woodcroft, B. J., Singleton, C. M., Boyd, J. A., Evans, P. N., Emerson, J. B., et al. (2018). Genome-centric view of carbon processing in thawing permafrost. Nature, 560 (7716), 49–54. 10.1038/s41586-018-0338-1

Yang, Y., et al. (2021). Deciphering factors driving soil microbial life-history strategies in restored grasslands. Environmental Microbiology Reports, 13 (2), 178–188. 10.1002/imt2.66

Zhang, Z., & Furman, A. (2021). Soil redox dynamics under dynamic hydrologic regimes - a review. Science of The Total Environment, 763, 143026. 10.1016/j.scitotenv.2020.143026

Zhou, Z., Tran, P. Q., Breister, A. M., Liu, Y., Kieft, K., Cowley, E. S., Karaoz, U., & Anantharaman, K. (2021). Metabolic: High-throughput profiling of microbial genomes for functional traits, biogeochemistry, and community-scale metabolic networks. Microbiome, 9 (1), 102. 10.1186/s40168-021-01213-8

