## Supplementary Information for "Metagenomic Insights Into Microbial Controls of Carbon Cycling in Alpine Soils"

**for**

### 1 Supplementary Figures

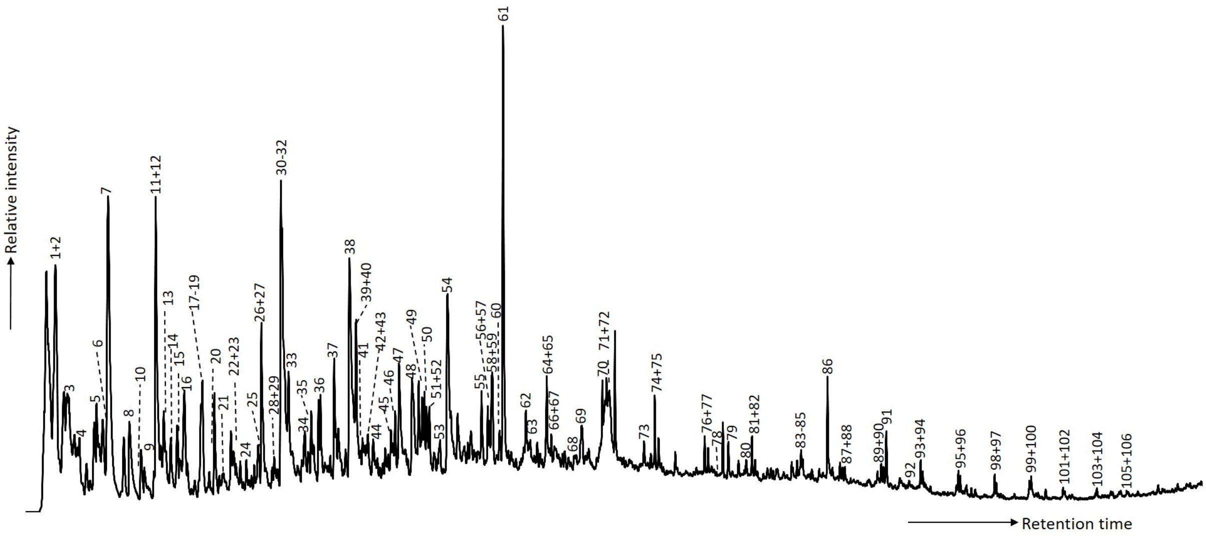


Figure S1: Example pyrogramm from pyrolysis gas chromatography–mass spectrometry analysis.

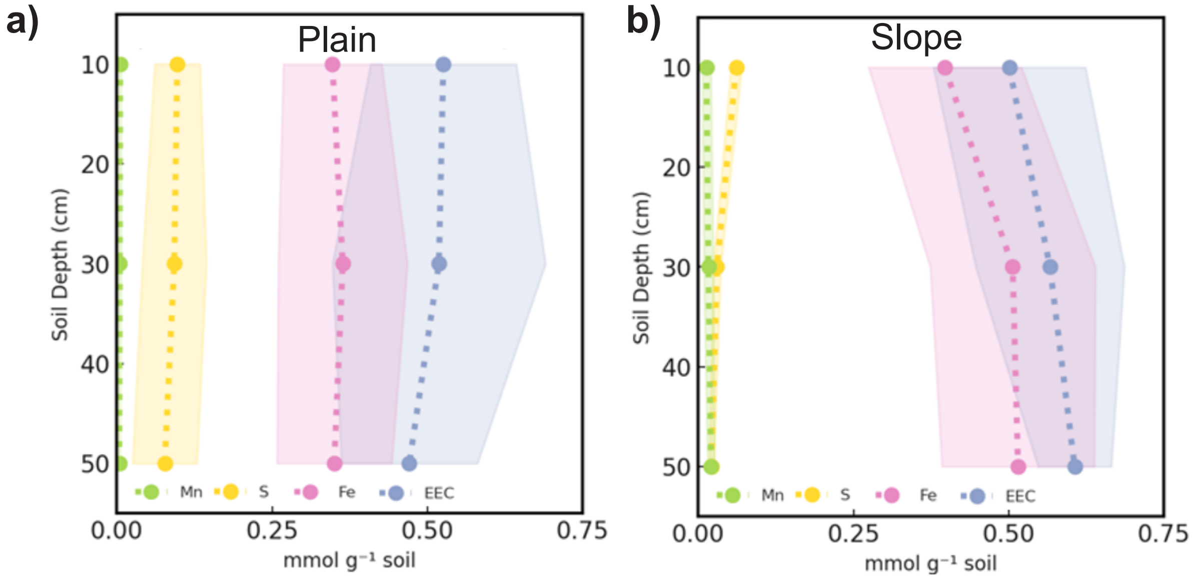


Figure S2: Electron exchanging capacity (EEC) and exchange capacities for iron, sulfur, and manganese. EEC values were determined from the sum of electron donating and accepting capacity. Element-specific exchange capacities were calculated from elemental concentrations, assuming one electron exchange per atom. Shaded areas represent the standard error of the mean. Each data point reflects the average of five soil samples.

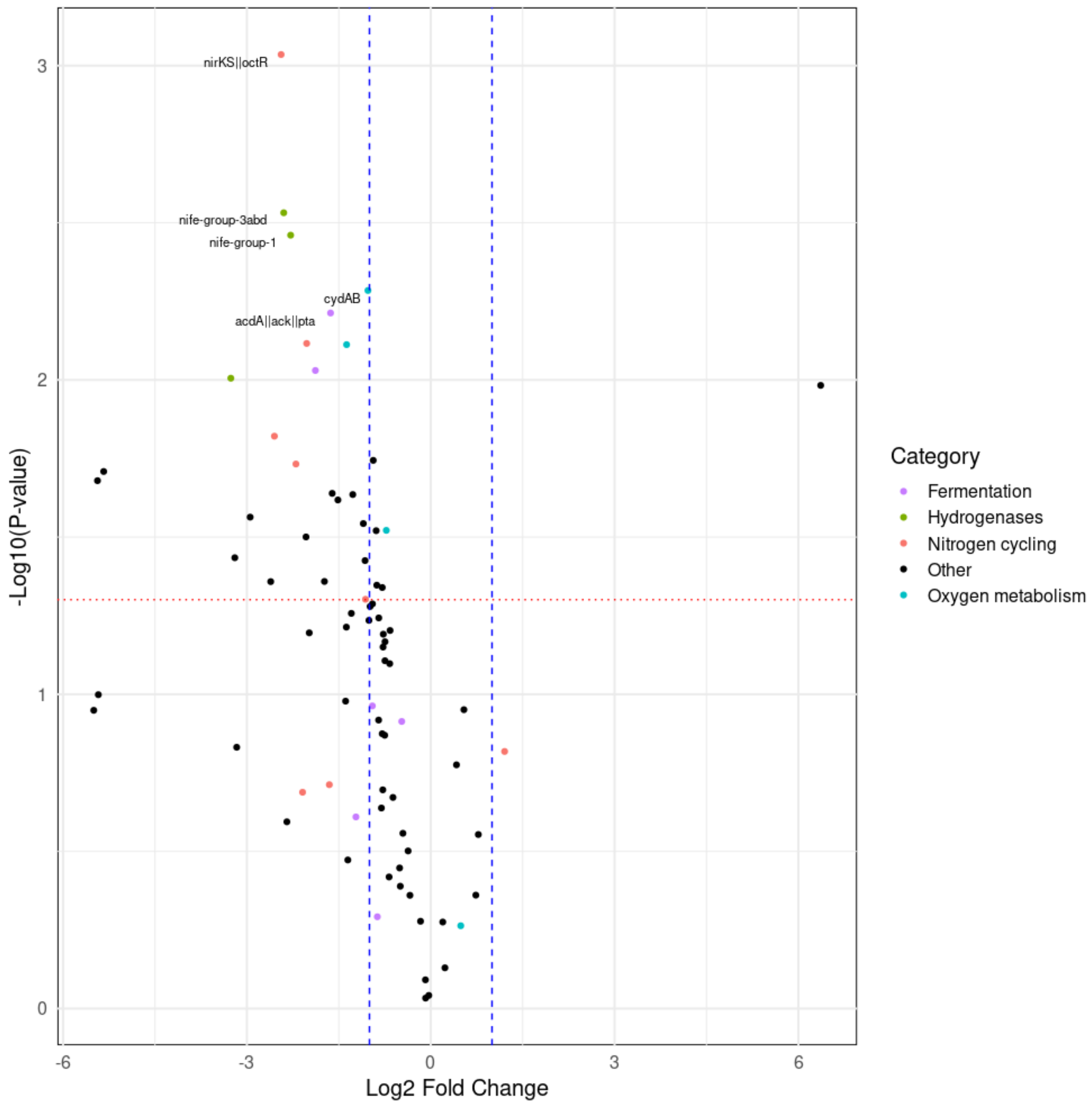


Figure S3: Volcano plot showing differential gene expression for metabolic pathways between plain (left) and slope soils (right).

#### 2 Supplementary Tables

Table S1: Soil organic carbon composition of plain and slope soils. Values represent the relative abundance of each compound class as a percentage of total SOC, determined by pyrolysis gas chromatography–mass spectrometry.

| ID | Catchment | Position | Depth | X | Y | Lipids | Aromatics | Lignins | Polysacch. | N compounds | Phenols |
|---|---|---|---|---|---|---|---|---|---|---|---|
| 1 | Binntal | Plain | 0–10 cm | 2664551 | 1136956 | 5.3 | 23.0 | 8.8 | 26.2 | 11.2 | 25.4 |
| 2 | Binntal | Plain | 10–30 cm | 2664551 | 1136956 | 5.9 | 21.2 | 10.4 | 21.7 | 11.6 | 29.1 |
| 3 | Binntal | Plain | 30–50 cm | 2664551 | 1136956 | 6.8 | 28.7 | 4.3 | 26.2 | 12.9 | 21.0 |
| 4 | Binntal | Plain | 0–10 cm | 2664545 | 1136922 | 4.3 | 19.9 | 7.7 | 23.7 | 10.4 | 34.1 |
| 5 | Binntal | Plain | 10–30 cm | 2664618 | 1136947 | 5.4 | 36.2 | 2.1 | 21.9 | 13.0 | 21.5 |
| 6 | Binntal | Plain | 30–50 cm | 2664618 | 1136947 | 32.1 | 24.4 | 4.4 | 14.8 | 10.6 | 13.7 |
| 7 | Binntal | Slope | 0–10 cm | 2664534 | 1137140 | 6.1 | 25.0 | 3.2 | 39.9 | 11.4 | 14.4 |
| 8 | Binntal | Slope | 10–30 cm | 2664534 | 1137140 | 4.6 | 39.9 | 0.9 | 32.0 | 12.4 | 10.2 |
| 9 | Binntal | Slope | 30–50 cm | 2664534 | 1137140 | 7.2 | 51.0 | 0.2 | 18.7 | 14.5 | 8.5 |
| 10 | Binntal | Slope | 0–10 cm | 2664594 | 1137136 | 5.5 | 15.7 | 9.0 | 46.1 | 11.2 | 12.5 |
| 11 | Binntal | Slope | 10–30 cm | 2664590 | 1137072 | 13.6 | 28.3 | 1.5 | 35.7 | 11.8 | 9.1 |
| 12 | Réchy | Plain | 10–30 cm | 2605594 | 1116454 | 11.8 | 27.5 | 4.4 | 17.9 | 12.9 | 25.5 |
| 13 | Réchy | Plain | 30–50 cm | 2605594 | 1116454 | 7.6 | 35.7 | 1.3 | 28.4 | 12.4 | 14.5 |
| 14 | Réchy | Plain | 0–10 cm | 2605565 | 1116589 | 7.1 | 25.6 | 9.0 | 18.4 | 14.5 | 25.5 |
| 15 | Réchy | Plain | 10–30 cm | 2605457 | 1116566 | 7.1 | 43.8 | 1.0 | 17.5 | 19.1 | 11.4 |
| 16 | Réchy | Slope | 0–10 cm | 2605229 | 1116612 | 4.0 | 20.9 | 15.0 | 25.7 | 13.0 | 21.5 |
| 17 | Réchy | Slope | 10–30 cm | 2605278 | 1116631 | 8.3 | 27.4 | 1.9 | 27.1 | 19.5 | 15.7 |
| 18 | Réchy | Slope | 30–50 cm | 2605278 | 1116631 | 5.2 | 27.9 | 0.8 | 29.6 | 20.6 | 15.8 |
| 19 | Réchy | Slope | 0–10 cm | 2605130 | 1116572 | 6.4 | 26.5 | 2.0 | 37.0 | 13.8 | 14.4 |
| 20 | Réchy | Slope | 10–30 cm | 2605273 | 1116750 | 4.4 | 37.4 | 0.3 | 27.7 | 19.9 | 10.2 |

Table S2: Environmental parameters used in NMDS ordination. Values represent soil pH, Ca content, and soil texture fractions measured at each sampling location (see Table S1).

| ID | pH | Clay (% v/v) | Silt (% v/v) | Sand (% v/v) | Ca (µg/g) |
|---|---|---|---|---|---|
| 1 | 7.12 | 0.99 | 34.59 | 64.42 | 52168.33 |
| 2 | 7.28 | 1.50 | 49.13 | 49.37 | 32066.74 |
| 3 | 6.91 | 0.76 | 25.83 | 73.41 | 45242.11 |
| 4 | 7.54 | 1.41 | 46.47 | 52.11 | 24799.30 |
| 5 | 7.51 | 0.76 | 30.94 | 68.30 | 13953.14 |
| 6 | 7.79 | 0.99 | 31.35 | 67.66 | 24174.14 |
| 7 | 4.85 | 2.92 | 33.81 | 63.27 | 7360.88 |
| 8 | 5.02 | 3.52 | 34.15 | 62.33 | 7098.25 |
| 9 | 5.22 | 4.95 | 41.37 | 53.69 | 6835.10 |
| 10 | 5.21 | 1.96 | 31.09 | 66.96 | 7492.29 |
| 11 | 5.02 | 3.39 | 38.72 | 57.88 | 6729.88 |
| 12 | 6.52 | 3.14 | 80.16 | 16.70 | 5291.61 |
| 13 | 6.63 | 3.97 | 74.20 | 21.83 | 4032.77 |
| 14 | 5.59 | 3.71 | 84.50 | 11.80 | 5006.22 |
| 15 | 6.32 | 3.13 | 72.04 | 24.83 | 3629.57 |
| 16 | 6.85 | 3.40 | 55.34 | 41.26 | 4279.37 |
| 17 | 6.89 | 4.87 | 63.37 | 31.76 | 5306.71 |
| 18 | 6.96 | 4.53 | 64.57 | 30.90 | 4996.69 |
| 19 | 5.23 | 5.35 | 59.27 | 35.38 | 2835.53 |
| 20 | 6.47 | 3.88 | 51.64 | 44.47 | 3028.28 |

Table S3: List of ppyrolysis gas chromatography–mass spectrometry moieties found in the studied soils, containing peak number (as related to Figure S1), retention time, compound class, molecular weight and masses used quantification. [a] ncc = nitrogen containing compounds

| **Pyrolysis moiety** | **Peak #** | **Retention time (min)** | **Compound class**[a] | **Molecular weight** | **Masses** |
|---|---|---|---|---|---|
| 2-methylfurnan | 1 | 2.4 | polysaccharides | 82 | 53+82 |
| acetic acid | 2 | 2.4 | polysaccharides | 60 | 60 |
| benzene | 3 | 3.0 | aromatics | 78 | 77+78 |
| (1H)-pyrrole, dimethyl | 4 | 3.4 | ncc | 96 | 95+96 |
| Pyridine | 5 | 4.2 | ncc | 79 | 52+79 |
| Pyrrole | 6 | 4.5 | ncc | 67 | 67 |
| toluene | 7 | 4.7 | aromatics | 92 | 92+91 |
| (2H)-furan-3-one | 8 | 5.7 | polysaccharides | 84 | 54+84 |
| 3 furaldehyde | 9 | 6.2 | polysaccharides | 96 | 95+96 |
| methylpyridine | 10 | 6.4 | ncc | 93 | 66+93 |
| cyclopenten-1-one | 11 | 6.8 | polysaccharides | 82 | 82+54 |
| Furfural | 12 | 6.9 | polysaccharides | 96 | 95+96 |
| methyl-1H-pyrrole | 13 | 7.2 | ncc | 81 | 80+81 |
| methyl-1H-pyrrole | 14 | 7.5 | ncc | 81 | 80+81 |
| C2 bezene (xylene) | 15 | 7.8 | aromatics | 106 | 106+91 |
| C2 bezene (xylene) | 16 | 8.1 | aromatics | 106 | 106+91 |
| styrene | 17 | 8.9 | aromatics | 104 | 104+78 |
| C2 bezene (xylene) | 18 | 9.0 | aromatics | 106 | 106+91 |
| c9 alkene | 19 | 9.0 | lipids | 126 | 55+69 |
| C9 alkane | 20 | 9.3 | lipids | 128 | 57+71 |
| acetylfuran | 21 | 9.7 | polysaccharides | 110 | 110+95 |

*Continue on the next page*

Table S3: (cont.)

| **Pyrolysis moiety** | **Peak #** | **Retention time (min)** | **Compound class**$^{a}$ | **Molecular weight** | **Masses** |
|---|---|---|---|---|---|
| 2hydroxy-2-cyclopenten-1-one | 22 | 10.3 | polysaccharides | 98 | 98+55 |
| dimethylpyridine | 23 | 10.5 | ncc | 107 | 106+107 |
| propylbenzene | 24 | 11.2 | aromatics | 120 | 120+91 |
| C3 benzene | 25 | 11.5 | aromatics | 120 | 105+120 |
| 5 methyl fufural | 26 | 11.6 | polysaccharides | 110 | 110+109 |
| C3 benzene | 27 | 11.7 | aromatics | 120 | 105+120 |
| C3 benzene | 28 | 12.1 | aromatics | 120 | 105+120 |
| benzonitrile | 29 | 12.3 | ncc | 103 | 76+103 |
| Phenol | 30 | 12.5 | phenols | 94 | 94+66 |
| c10 alkene | 31 | 12.5 | lipids | 140 | 55+69 |
| C3 benzene | 32 | 12.6 | aromatics | 120 | 105+120 |
| C10 alkane | 33 | 12.9 | lipids | 142 | 57+71 |
| C3 benzene | 34 | 13.6 | aromatics | 120 | 105+120 |
| 3-hydroxy-2-methyl-2-cyclopenten-1-one | 35 | 13.9 | polysaccharides | 112 | 112 |
| Indene | 36 | 14.3 | aromatics | 116 | 116+115 |
| methylphenol | 37 | 14.9 | phenols | 108 | 107+108 |
| methylphenol | 38 | 15.6 | phenols | 108 | 107+108 |
| 4-methoxyphenol (guaicol) | 39 | 15.9 | lignins | 124 | 109+124 |
| c11 alkene | 40 | 15.9 | lipids | 154 | 55+69 |
| C11 alkane | 41 | 16.2 | lipids | 156 | 57+71 |
| methylbenzofuran | 42 | 16.3 | polysaccharides | 132 | 132+131 |

*Continue on the next page*

Table S3: (cont.)

| **Pyrolysis moiety** | **Peak #** | **Retention time (min)** | **Compound class**$^{a}$ | **Molecular weight** | **Masses** |
|---|---|---|---|---|---|
| methylbenzofuran | 43 | 16.4 | polysaccharides | 132 | 132+131 |
| maltol | 44 | 16.8 | polysaccharides | 126 | 126 |
| benzylnitrile | 45 | 17.5 | ncc | 117 | 90+117 |
| 3methyl 1H-indene | 46 | 17.6 | aromatics | 130 | 130+115 |
| dimethyl/ethylphenol | 47 | 17.8 | phenols | 122 | 107+122 |
| dimethyl/ethylphenol | 48 | 18.4 | phenols | 122 | 107+122 |
| naphthalene | 49 | 18.7 | aromatics | 128 | 128 |
| c12 alkene | 50 | 19.0 | lipids | 168 | 55+69 |
| 4-methylguiaacol (creosol) | 51 | 19.1 | lignins | 138 | 123+138 |
| C12 alkane | 52 | 19.2 | lipids | 170 | 57+71 |
| 4,7-dimethylbenzofuran | 53 | 19.7 | polysaccharides | 146 | 145+146 |
| 4-vinylphenol | 54 | 20.1 | phenols | 120 | 120+91 |
| 4-ethylguaiacol | 55 | 21.5 | lignins | 152 | 137+152 |
| c13 alkene | 56 | 21.8 | lipids | 182 | 55+69 |
| methyl naphthalene | 57 | 21.9 | aromatics | 142 | 142+141 |
| Indole | 58 | 22.0 | ncc | 117 | 90+117 |
| C13 alkane | 59 | 22.0 | lipids | 184 | 57+71 |
| methyl naphthalene | 60 | 22.4 | aromatics | 142 | 142+141 |
| 4-vinylguaiacol | 61 | 22.5 | lignins | 150 | 135+150 |
| 2,6-dimethoxyphenol (syringol) | 62 | 23.6 | lignins | 154 | 139+154 |
| 4-Propenylguaiacol | 63 | 23.7 | lignins | 164 | 164+149 |

*Continue on the next page*

Table S3: (cont.)

| **Pyrolysis moiety** | **Peak #** | **Retention time (min)** | **Compound class**[a] | **Molecular weight** | **Masses** |
|---|---|---|---|---|---|
| methyl indole | 64 | 24.5 | ncc | 131 | 131+130 |
| c14 alkene | 65 | 24.5 | lipids | 196 | 55+69 |
| C14 alkane | 66 | 24.7 | lipids | 198 | 57+71 |
| 4-formylguaicol (Vanillin) | 67 | 24.9 | lignins | 152 | 151+152 |
| 4-methylsyringol | 68 | 26.0 | lignins | 168 | 153+168 |
| trans-4-(2-propenyl)guaiacol (eugenol) | 69 | 26.0 | lignins | 164 | 164+149 |
| c15 alkene | 70 | 27.0 | lipids | 210 | 55+69 |
| levoglucosan | 71 | 27.3 | polysaccharides | 162 | 73+60 |
| C15 alkane | 72 | 27.2 | lipids | 212 | 57+71 |
| 4-vinylsyringol | 73 | 28.9 | lignins | 180 | 165+180 |
| c16 alkene | 74 | 29.3 | lipids | 224 | 55+69 |
| C16 alkane | 75 | 29.5 | lipids | 226 | 57+71 |
| c17 alkene | 76 | 31.6 | lipids | 238 | 55+69 |
| C17 alkane | 77 | 31.8 | lipids | 240 | 57+71 |
| diketodipyrrole | 78 | 32.0 | NCC | 186 | 93+186 |
| 4-acetylsyringol | 79 | 32.7 | lignins | 196 | 181+196 |
| phenanthrene | 80 | 33.5 | aromatics | 178 | 178 |
| c18 alkene | 81 | 33.7 | lipids | 252 | 55+69 |
| C18 alkane | 82 | 33.9 | lipids | 254 | 57+71 |
| c19 alkene | 83 | 35.8 | lipids | 266 | 55+69 |
| C19 alkane | 84 | 35.9 | lipids | 268 | 57+71 |

*Continue on the next page*

Table S3: (cont.)

| **Pyrolysis moiety** | **Peak #** | **Retention time (min)** | **Compound class**$^{a}$ | **Molecular weight** | **Masses** |
|---|---|---|---|---|---|
| C17 methylketone | 85 | 36.0 | lipids | 254 | 58+59 |
| C16 fatty acid | 86 | 37.1 | lipids | 256 | 60+73 |
| c20 alkene | 87 | 37.7 | lipids | 280 | 55+69 |
| C20 alkane | 88 | 37.8 | lipids | 282 | 57+71 |
| C21 alkene | 89 | 39.5 | lipids | 294 | 55+69 |
| C21 alkane | 90 | 39.7 | lipids | 296 | 57+71 |
| C19 methylketone | 91 | 39.8 | lipids | 282 | 58+59 |
| C18 fatty acid | 92 | 40.8 | lipids | 284 | 60+73 |
| C22 alkene | 93 | 41.3 | lipids | 308 | 55+69 |
| C22 alkane | 94 | 41.4 | lipids | 310 | 57+71 |
| C23 alkene | 95 | 43.0 | lipids | 322 | 55+69 |
| C23 alkane | 96 | 43.1 | lipids | 324 | 57+71 |
| C24 alkene | 97 | 44.7 | lipids | 336 | 55+69 |
| C24 alkane | 98 | 44.8 | lipids | 338 | 57+71 |
| C25 alkene | 99 | 46.2 | lipids | 350 | 55+69 |
| C25 alkane | 100 | 46.3 | lipids | 352 | 57+71 |
| C26 alkene | 101 | 47.8 | lipids | 364 | 55+69 |
| C26 alkane | 102 | 47.8 | lipids | 366 | 57+71 |
| C27 alkene | 103 | 49.2 | lipids | 378 | 55+69 |
| C27 alkane | 104 | 49.3 | lipids | 380 | 57+71 |
| C28 alkane | 105 | 50.7 | lipids | 394 | 57+71 |
| C29 alkane | 106 | 52.0 | lipids | 408 | 57+71 |
